# Liquid-Liquid Phase Separation-mediated formation of amyloid fibrils from DcpS scavenger enzymes

**DOI:** 10.1101/2025.09.24.678264

**Authors:** Aleksandra Ferenc-Mrozek, Maria Winiewska-Szajewska, Hanna Nieznańska, Wojciech Dzwolak, Maciej Łukaszewicz

**Affiliations:** Division of Biophysics, Faculty of Physics, University of Warsaw, Pasteura 5, 02-093 Warsaw, Poland; Institute of Biochemistry and Biophysics PAS, Pawinskiego 5a, 02-106, Warsaw, Poland; Laboratory of Electron Microscopy, Nencki Institute of Experimental Biology PAS, Pasteura 3, 02-093 Warsaw, Poland; Faculty of Chemistry, Biological and Chemical Research Centre, University of Warsaw, Pasteura 1, 02-093 Warsaw, Poland

**Author notes:** Corresponding authors: Maciej Lukaszewicz, Wojciech Dzwolak.

**Keywords:** DcpS, decapping, LLPS, amyloid fibrils

## Abstract

Decapping Scavenger (DcpS) enzyme was initially identified by its ability to hydrolyze the cap structure resulting from mRNA decay. Human DcpS is an established target for acute myeloid leukemia (AML) and hepatic metastasis. Recently, the protein has been linked to neuronal development regulation and implicated in certain developmental neurological disorders. Here we demonstrate for the first time that DcpS undergoes misfolding in vitro, leading to the formation of amyloid-like fibrils. Fibrillization was observed for human and nematode (C. elegans) DcpS using transmission electron microscope (TEM) imaging, Thioflavin T (ThT) fluorescence assay, Fourier-transform infrared (FT-IR) spectroscopy, circular dichroism (CD) spectroscopy, differential scanning fluorimetry (DSF), and dynamic light scattering (DLS). Additionally, the DcpS^INS15^ insertional mutant linked to the Al-Raquad syndrome, exhibited accelerated fibril aggregation kinetics compared to the wild type protein. Moreover, we show that the DcpS species investigated in this study undergo liquid-liquid phase separation (LLPS) prior to amyloid-formation. We propose that the LLPS phase transition underlies the intricate kinetics (e.g. lack of a clearly-resolved lag phase) of the misfolding process. As the physiological implications of the here-reported propensity of DcpS to lose its biological function through the coupled LLPS-fibrillization transition remain to be elucidated, this work lays the groundwork for further studies on this phenomenon.

## Introduction

Most proteins have a well-defined, stable three-dimensional structure under close-to-physiological conditions. Stress factors like elevated temperature^1^, pressure^2^, non-physiological pH^3^ or presence of chaotropic cosolutes and cosolvents^4^ perturb protein’s native structure. Depending on the type of perturbation the unfolding process can acquire different characteristics^5^. Destabilized proteins can undergo transition into aggregates either directly from the native state or via a transient intermediate state^6^. Those fully-, or partially unfolded intermediates (e.g. molten-globule-like states^7^) serve as aggregation precursors^8^. Many proteins contain one or more specific short sequence segments consisting of 5-15 hydrophobic amino acids, known as aggregation-prone regions (APRs)^9^. In the protein’s native state, APRs are typically buried within a hydrophobic core but, under unfolding conditions, the likelihood of APR exposure to the solvent increases. Exposure of these segments facilitates self-association between partially unfolded molecules, thereby triggering aggregation^10^. In general, aggregation is a highly complex process, yielding productss varying in morphology, ranging from soluble oligomers to insoluble amorphous aggregates and amyloid fibrils^11^. A characteristic feature of amyloid aggregates is their hierarchical structural organization, wherein fibrils are formed through the lateral association of protofilaments assembled from stacked β-sheets^12^. The formation of amyloid fibrils is closely associated with both biological functions (e.g. formation of functional amyloids) and human diseases^13^.

Amyloid aggregation is a hallmark of numerous degenerative disorders, including Alzheimer’s disease (AD) and other tauopathies, Parkinson’s disease, type II diabetes^14^, and systemic amyloidosis e.g. immunoglobulin light chain (AL) amyloidosis^15^. In humans, at least 42 proteins have been identified in their amyloid, disease-associated forms^16^, with additional candidates under investigation^17^. Several mechanisms of amyloid formation have been proposed^18^, with the most widely accepted being the nucleation-dependent polymerization model, characterized by a sigmoidal aggregation kinetic curve. This model proposes that amyloid fibril formation occurs through an initial lag phase, followed by elongation of fibrils ^19–22^. Other mechanisms (e.g. nucleation-independent^23^) have also been proposed. In recent years, increasing attention has been given to the role of liquid-liquid phase separation (LLPS) in pathological protein aggregation. LLPS is an intriguing phenomenon observed in cellular environments, leading to the formation of membraneless organelles or biomolecular condensates involved in various cellular processes, including signaling, transcription, RNA processing and protection under stress conditions^24–28^. While the formation of droplets via the LLPS pathway is typically reversible, if proteins within the droplets are subjected to persistent stress conditions, or if the LLPS-induced increased protein concentration is coupled with structure-destabilizing mutations in the protein sequence, the occurrence of droplets may facilitate formation of insoluble aggregates within^29–31^. Emerging evidence suggests that some amyloidogenic proteins implicated in various diseases, including neurodegenerative disorders such as amyotrophic lateral sclerosis (ALS) and frontotemporal dementia (FTD) or cancer, can form droplets prior to amyloid fibrils^32–35^.

Here, we investigated the propensity of the decapping scavenger enzyme (DcpS) to form fibrillar aggregates under thermal stress conditions and at the physiological temperature (37°C). DcpS belongs to the superfamily of HIT (Histidine Triad) proteins, which includes hydrolases and nucleotide transferases^36,37^. DcpS proteins are found in eukaryotes of varying complexity, including humans, nematodes and yeast^38,39,^. Screening experiments have confirmed that the gene encoding human DcpS is one of the factors that determine cell viability^40^. The primary recognized function of DcpS is its role in mRNA turnover^41^. It hydrolyzes dinucleotide cap structures or short capped oligonucleotides resulting from the processive exosome mRNA degradation in the 3’ to 5’ direction^38,42^. Independent of its catalytic activity, DcpS also functions as a transcript-specific modulator^43^ involved in the pre-mRNA splicing ^44^ and regulation of miRNA turnover^45^. Importantly, potential relevance of DcpS in certain clinical conditions has been reported. Human DcpS has been considered as a diagnostic biomarker associated with the colorectal cancer^46^ and as a therapeutic target for hepatic metastasis in uveal melanoma^47^ and acute myeloid leukemia (AML)^48^. Since 2015, several mutations in the *DcpS* gene have been linked to certain intellectual disabilities^49,50^. Clinically diagnosed disorders affecting the heart, joints, and muscles, as well as minor deformations within the craniofacial region, have been classified as Al-Raqad Syndrome (ARS)^51^. Additionally, magnetic resonance imaging of a patient’s brain has revealed progressive demyelination of the central and peripheral nervous system^52^. Studies on cells from patients with DcpS enzyme dysfunction have shown impaired differentiation and growth of neurites^53^. This highlights the crucial role of DcpS activity in neuronal development and suggests that impaired mRNA metabolism may be the underlying cause of neurological diseases. It appears that diverse physical and intellectual anomalies in individuals with ARS are determined by various types and locations of mutations in DcpS^51^.

There is limited information on the stability and aggregation-propensity of DcpS proteins in aqueous solutions, except for the melting temperature (T_m_) for human and nematode DcpS, obtained via dye-based differential scanning fluorimetry (DSF)^54^. Here, we conducted a more detailed study using biophysical methods to investigate the structural changes induced by thermal stress on human and *C. elegans* DcpS (CeDcpS). Using circular dichroism spectroscopy, we found that both proteins undergo irreversible aggregation, forming β-sheet-rich aggregates with rising temperature. These aggregates were further examined using FT-IR and TEM, confirming their morphology and structural features characteristic of amyloid fibrils. To monitor real-time aggregation at human physiological temperature, we employed a ThT binding assay. Our study also takes into account for the first time a mutated form of human DcpS, carrying the 15 amino acid insertion IKVSGWNVLISGHPA (DcpS^INS15^). Functional studies have demonstrated that the enzymatic activity of DcpS in patient-derived samples was significantly reduced, confirming the pathogenic nature of this mutation, which manifests in mental and physical disabilities^49^. Notably, the absence of a lag phase in fibril growth and the lack of a seeding were intriguing observations. We propose that this phenomenon may be explained by the occurrence of liquid-liquid phase separation, which is demonstrated here for DcpS.

## Results

### 1. Aggregation prone regions of DcpS proteins

Initially, we employed the WALTZ algorithm^55^ to identify putative amyloidogenic sequences in DcpS (**Figure 1**, **Supplementary Table 1**). For the human native DcpS enzyme, WALTZ identified two potentially amyloidogenic amino acid regions within the C-terminal domain of the protein: ^173^IQWVYNI^179^ (segment no.1) and ^199^GFVLIP^204^ (segment no.2). The mutant human protein (DcpS^INS15^) has an additional region compared to the unmutated form, located within the 15-amino acid insertion: ^218^WNVLIS^223^*. C. elegans* DcpS homologue of human protein (which shares 41% of the amino acid sequence identity with human DcpS) appears to have also two segments in its sequence that can potentially contribute to amyloid formation, likewise located within C-terminal domain: ^147^LNWVYN^152^ (segment no.1) and ^187^LENLYVLAI^195^ (segment no.2). The segment no.1 overlaps in the amino acid sequence alignment of both proteins and is located on the surface of the resolved human DcpS structure (**Figure 1C**), and on the surface of the modeled *C.elegans* DcpS structure (**Supplementary Figure 1B**).

**Figure 1.**
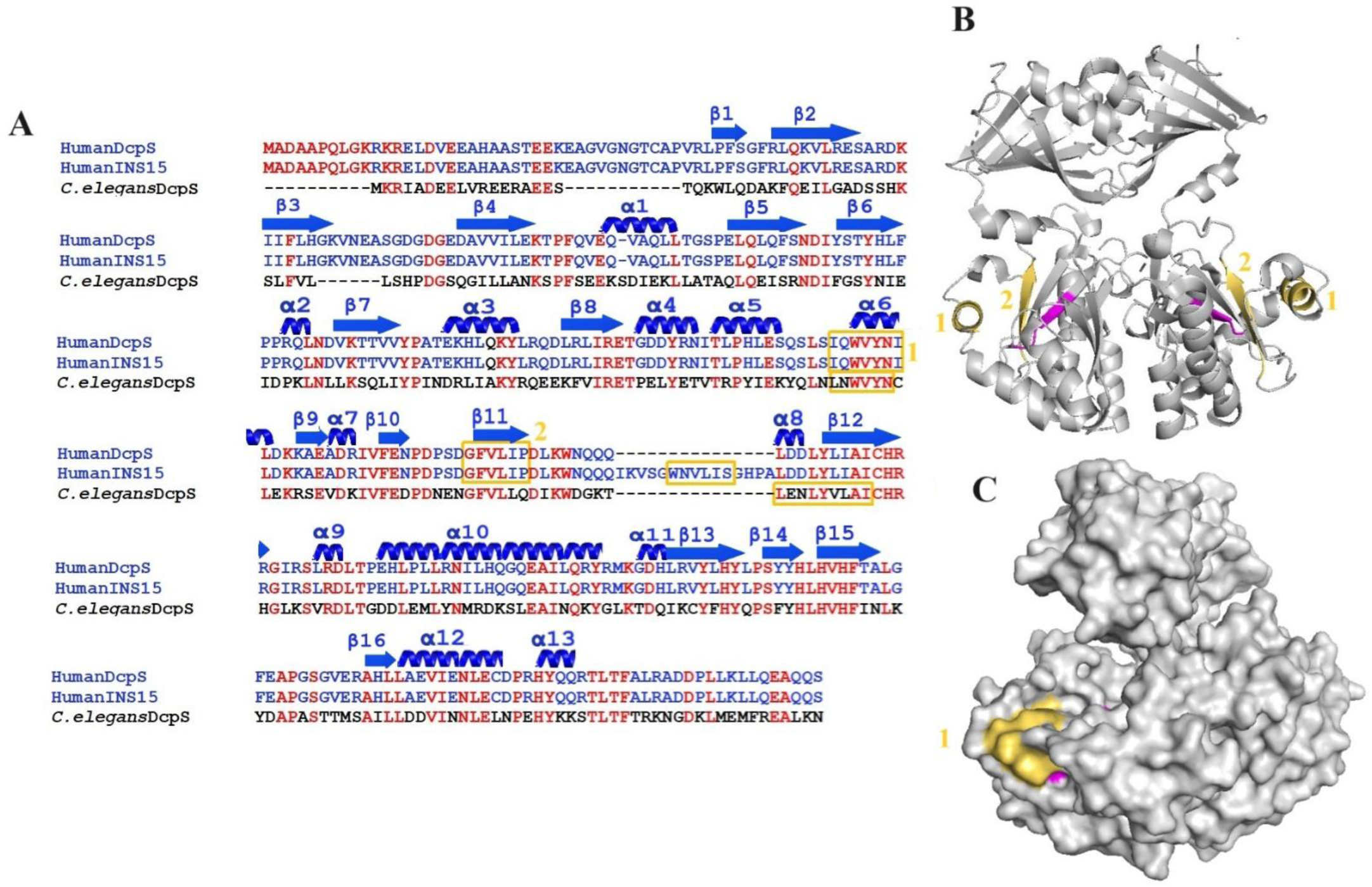
Prediction of aggregation-prone regions in DcpS proteins. **A** - Alignment of human DcpS, mutant human DcpS (Human^INS15^) and *C.elegans* DcpS, with secondary structures location (α-helix and β-sheet) for known crystal structure of human DcpS. Amino acids labeled in red in the sequence are identical for all three proteins. Aggregation-prone segments are highlighted with a yellow box. *C.elegans* DcpS and human DcpS possess two APRs, while the mutant human protein has an additional region within the insertion of 15 amino acids. **B, C** - Crystal structure of human DcpS were drawn based on PDB:1XML. Human DcpS APRs are colored yellow and catalytic centers (HIT motif) colored by magenta. APRs prediction was conducted using WALTZ. Structural images were prepared using PyMol.

### 2. Thermal denaturation of DcpS results in β-sheet aggregates

Temperature-denaturated proteins often form β-sheet-rich amyloid-like aggregates^56,57^. Initially, we decided to simultaneously monitor the thermal unfolding transition of DcpS proteins and aggregation in real-time using nano-differential scanning fluorimetry and dynamic light scattering. This approach relies on the change in intrinsic tryptophan fluorescence and aggregation-induced scattering of light by particles^58^. In **Figure 2**, the fluorescence intensity ratio at 350 and 330 nm as a function of temperature (**Figure 2 A, E**) is presented, accompanied by the light scattering thermograms (**Figure 2 B, F**). The values obtained for the folding state transition and aggregation temperatures were nearly identical for the particular DcpS protein: Tm∼57°C and Tagg∼57°C for human DcpS, and Tm∼47°C and Tagg∼48°C for *C.elegans* DcpS, (which is in the range of the previously reported Tm values based on DSF with SYPRO dye^54^). The aggregation propensity of DcpS in response to increasing temperature was further confirmed by differential scanning calorimetry (DSC), where the resulting thermograms showed both endo- and exothermic effects (**Supplementary Figure 2**). Protein denaturation is generally an endothermic process, while exothermic effects typically indicate protein aggregation in solution. Because of its irreversibility, the aggregation that occurs during the protein unfolding precludes deriving quantitative thermodynamic data from such a process^59^.

**Figure 2.**
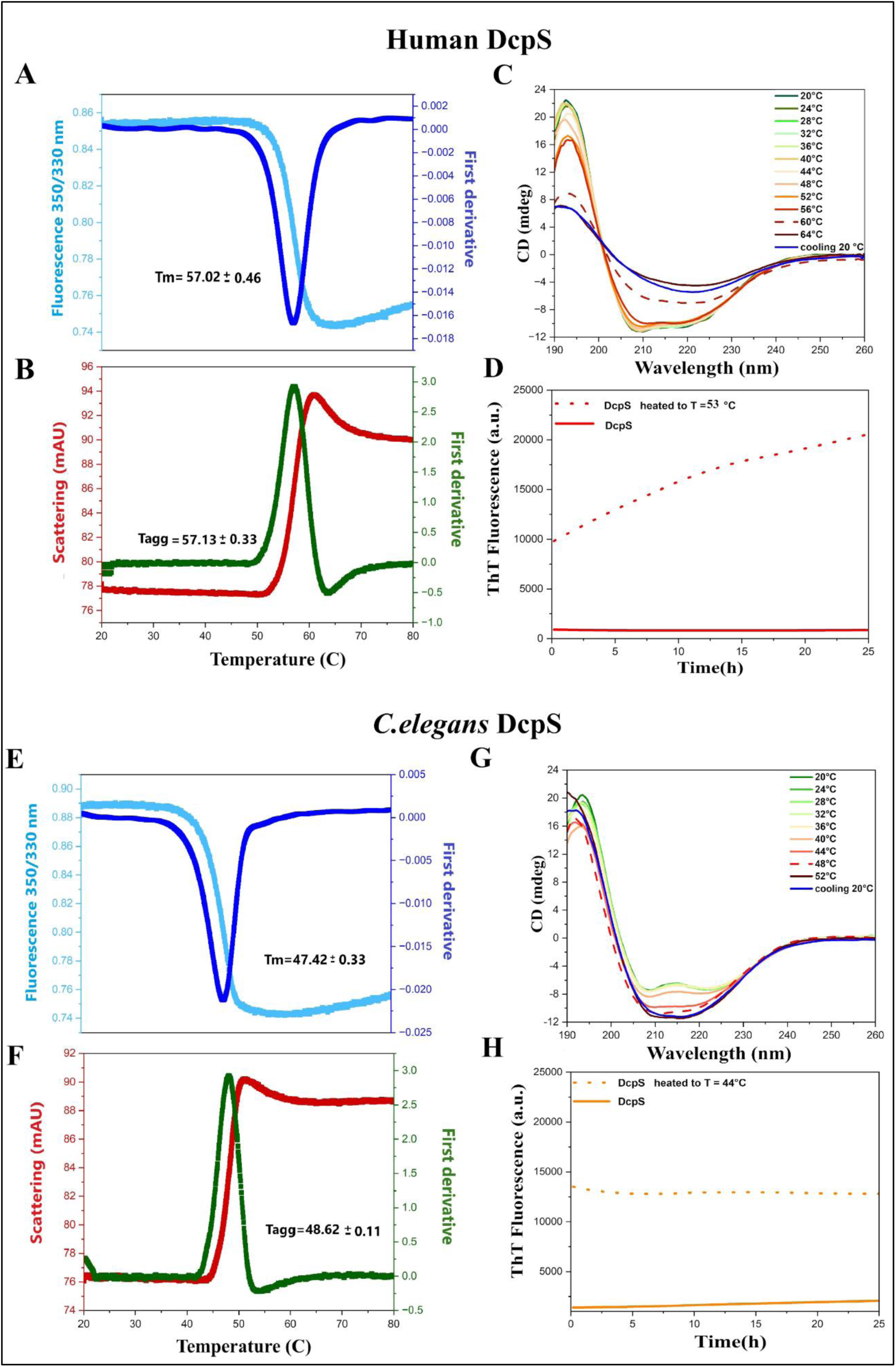
Thermal denaturation of human and *C.elegans* DcpS monitored by nanoDSF, DLS, and far-UV CD spectroscopy. **A,E-** Comparison of thermal stability of DcpS proteins by nanoDSF in temperature range 20°C – 80°C. Thermogram trajectories correspond to fluorescence ratio (Fluorescence intensity at 350nm/Fluorescence intensity at 330nm) -light blue, and it first derivative - dark blue. The temperature at which melting transition (Tm) occurs is indicated. Heating rate is 0.5°C/min. **B,F –** Scattering profile (red) in temperature range 20°C – 80°C. Calculated first derivative (green) corresponds to aggregation temperature. Heating rate is 0.5°C/min. **C, G** – Overlay of smoothed far-UV CD spectra obtained at temperatures ranging from 20°C to 64°C (for human DcpS) or to 52°C (for *C.elegans* DcpS) that is 4°C above the Tm temperature derived from nanoDSF (dotted line). Heating rate is 1°C/2min. In each case, green line is for the far-UV CD spectrum obtained at 20°C and the series of rainbow-colored lines are spectra plots obtained at 2°C intervals. Arrows indicate the direction of spectra development. In dark blue shown are the spectra obtained after cooling at 20°C for 12 hours of samples of human DcpS and *C.elegans* DcpS heated to 56°C and 44°C, respectively. **D, H -** Aggregation kinetics of *C.elegans* and human DcpS monitored by ThT fluorescence assay. Dotted lines show results obtained after heating up the DcpS sample to temperature 4°C below the nanoDSF-derived Tm, followed by incubation with Thioflavin T at 25°C. The solid lines represent fluorescence signal obtained for proteins not exposed to the raised temperature prior to start of the ThT assay.

In the DSF, DLS and ThT experiments - protein concentration was 0.5mg/ml in 50mM TRIS, 150mM NaCl, 0.2mM TCEP, pH 7.5. In the far-UV CD experiments – 0.2mg/ml DcpS was in 15mM TRIS, 80mM NaClO4, 0.2mM TCEP, pH 7.5 In order to gain a more detailed insight into the structural transition between the folded and unfolded states, as well as the aggregation process, we recorded far-UV CD spectra in the temperature range from 20°C to 64°C for human DcpS and from 20°C to 52°C for *C.elegans* DcpS, with a gradual increase at 1°C/2min intervals (**Figure 2 C, G**). At 20°C, the CD spectra of both DcpS proteins exhibited two distinct minima at 208nm and 222nm, indicative of α-helix secondary structure. The ellipticity for both the studied DcpS species vanished at temperature near the apparent Tm (dashed lines) (**Figure 2 C, G**), followed by further changes in measured CD signal. Attempts to renaturate the proteins by cooling to 20°C over a 12-hour period (following the heating to 64°C for human DcpS or 52°C for *C.elegans* DcpS) were unsuccessful. Also, cooling of once heated samples is not accompanied by a reversal of temperature-induced changes in the far-UV CD spectra, indicating that thermal denaturation of *C.elegans* and human DcpS is irreversible. Analysis of protein secondary structure content as a function of temperature based on CD measurements (using the BeStSel server)^60^ is summarized in **Supplementary Figure 3**. For human DcpS, the ratio of secondary structures remained constant up to 40°C. Above this temperature, the β-sheet content gradually increased while the α-helical content decreased (**Supplementary Figure 3A**). Noticeable changes in the secondary structure composition were observed around the melting temperature, as assessed using nanoDSF: from 56°C onwards, the β-sheet content began to dominate over α-helices, as reflected in the corresponding CD spectra, which showed a gradual decrease in ellipticity at 208nm and 222nm, with the minimum at 217nm, characteristic of β-sheet structures, becoming gradually more pronounced above 56°C (**Figure 2C**). In contrast, CD spectra of *C.elegans* DcpS showed a gradual decrease in the CD signal at 208nm and 222nm (**Figure 2G**), suggesting either a conformational changes of protein or secondary structure rearrangement as a function of temperature. The calculated secondary structure content remained unchanged up to 44°C, but above this temperature point the β-sheet fraction increased significantly (**Supplementary Figure 3B**). It is important to note that protein aggregation and subsequent sedimentation of aggregates may lead to a reduction in the actual concentration of protein in the optical pathway during a CD measurement. The secondary structure contents were calculated, however, under the assumption of an unchanged protein concentration.

The heat-induced aggregation process of hDcpS and CeDcpS was investigated further using a thioflavin T (ThT) fluorescence assay. Upon binding to stacked β-sheets, as present in amyloid fibrils, the quantum yield of ThT fluorescence increases by 2-3 orders of magnitude which is the basis of the dye’s diagnostic applications^61^. In **Figure 2D** and **2H**, ThT fluorescence intensity was monitored for 24 hours at room temperature following pre-heating of protein samples (at the heating rate of 1°C/2min), to the onset temperature of scattering curves (44°C for *C. elegans* DcpS and 53°C for human DcpS), **Figure 2 B, F**. Surprisingly, the initial ThT fluorescence of the partially denatured samples was significantly higher (by at least 20-fold) compared to the non-heated DcpS control. For human DcpS, ThT fluorescence further increased during incubation, whereas for *C. elegans* DcpS, it remained constant. This outcome suggests that DcpS proteins form ThT-positive aggregates, consistent with the findings from far-UV CD analysis.

### 3. Visualization and analysis of the self-assembly of DcpS-derived fibrils

The aggregation kinetics of DcpS were further analyzed at a constant temperature over an extended period (up to ∼250 hours) using the ThT assay. A temperature of 37°C (human physiological temperature) was chosen, which is lower than the T_m_ and T_agg_ for both explored DcpS. While *C. elegans* nematode naturally functions at lower temperatures (20-30°C), we opted to standardize the experimental conditions by maintaining the uniform incubation temperature. As shown in **Figure 3A and 3D**, the intensity of ThT fluorescence for DcpS proteins increased over incubation time and was strongly dependent on the used protein concentration (similarly to e.g. insulin^62^, amyloid-β^63^ or light chains of immunoglobulin^64^ - known for amyloids formation – where the ThT fluorescence intensity and the amount of formed amyloid fibrils correlates with the applied initial protein concentration). For CeDcpS, the ThT fluorescence reached plateau within approximately 24 hours, with a *T_50_* value (that corresponds to the time required to reach 50 % of the ThT maximum fluorescence intensity in a kinetic assay^65^) of ∼11-12 hours, independent of protein concentration. For human DcpS, the maximum ThT signal was reached after 250 hours or more, with *T_50_ ≥* 100 hours (at 1mg/ml and 0.5 mg/ml concentrations). The curves obtained showing the kinetics of human and nematode DcpS aggregation at 37°C point to the absence of a detectable lag phase, regardless of the protein concentration.

**Figure 3.**
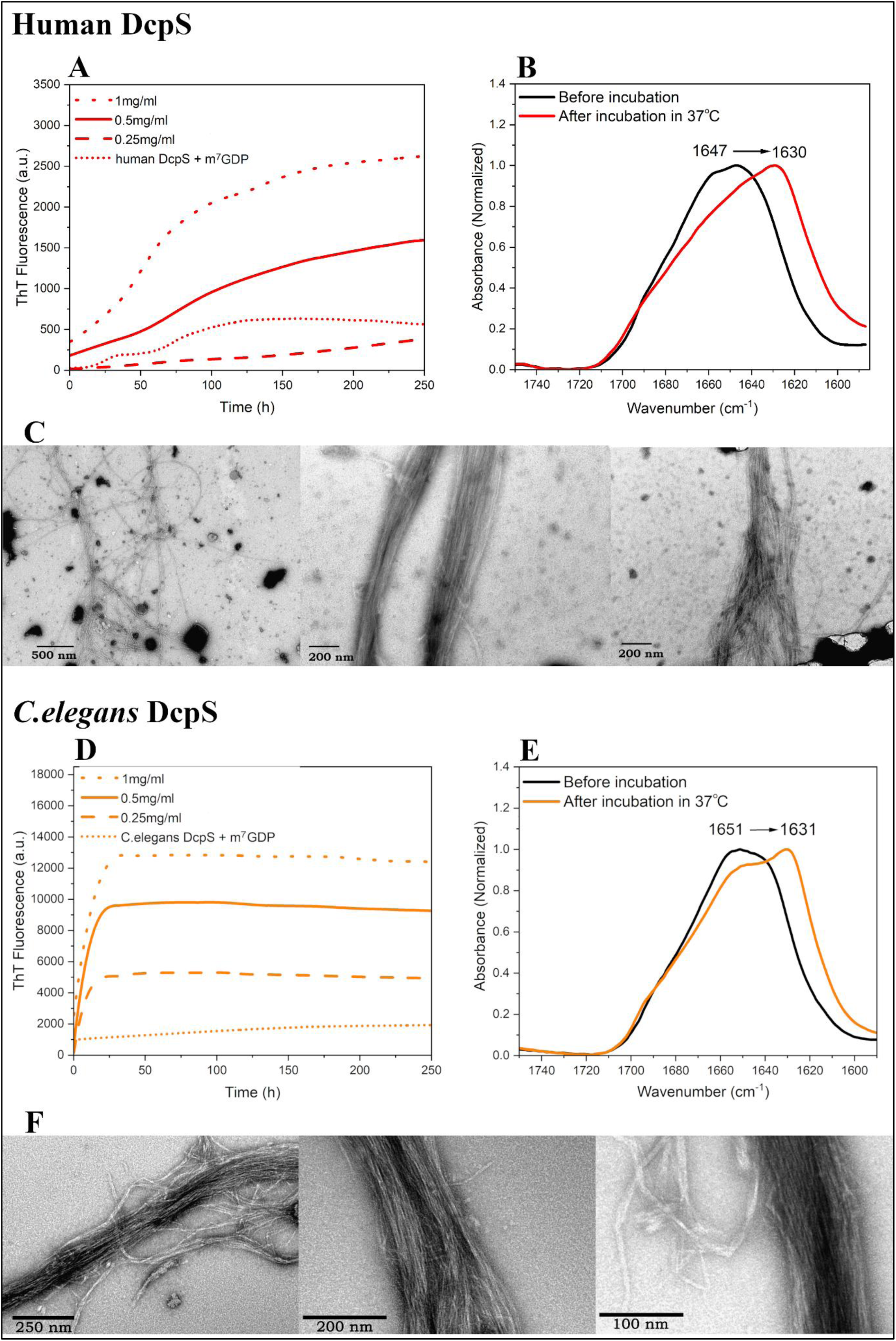
Amyloid-like fibril formation by human and *C.elegans* DcpS proteins. **A, D** - Kinetics of DcpS protein fibril formation probed by ThT fluorescence. Experiments were performed at 37°C for over 10 days of incubation for a range of DcpS concentrations: 0.25mg/ml (dashed line), 0.5mg/ml (solid line) and 1mg/ml (dotted line). No detectable lag phase is seen regardless of used protein concentration. Saturation of the ThT fluorescence signal and the end-state of fibril assembly is achieved faster for higher human DcpS concentrations. *C.elegans* DcpS fibrillization reaches plateau in almost the same time regardless of the concentration used. Data of ThT assay in the presence of DcpS ligand (m^7^GDP) at molar ratio 1:20 (ligand:protein), and at protein concentration 0.5mg/ml, is presented also (short dot line). DcpS was incubated with ligand for 30 minutes prior to ThT measurements. Representative set of data of three independent experiments is shown. **C, F** – TEM images of DcpS fibrillar aggregates, acquired after 10 days of incubation at 37°C. In the case of *C.elegans* DcpS, samples were sonicated before uranyl acetate staining and TEM imaging. Scale bares are indicated on images. Three representative micrographs are shown for each protein. **B, E** – FT-IR spectra acquired for human and *C.elegans* DcpS samples before and after 10 days incubation at 37 °C. Protein concentration at the beginning of experiment was 0.5mg/ml.

Tendency to amyloid aggregation can be limited by selective binding of ligands, thereby inhibiting the global process of unfolding under native or stressful conditions^66^. The m^7^GDP nucleotide is a well-known ligand that binds strongly to DcpS proteins^67^ and that it leads to the thermal stabilization of DcpS *in vitro*^54^. To investigate whether m^7^GDP could inhibit DcpS aggregation, we performed ThT assay in its presence. (**Figure 3A** and **D**). At a protein-to-ligand ratio of 1:20, the ThT fluorescence intensity for both human and nematode DcpS (at 0.5mg/ml concentration) was reduced approximately 5-6 fold compared to the sample without the ligand. Furthermore, the signal remains relatively stable even after nearly 250 hours of incubation, confirming that m^7^GDP binding significantly impedes the DcpS aggregation process. This inhibition is likely due to reduction in conformational fluctuations upon m^7^GDP binding. As the ligand stabilizes the population of native-like protein molecules, the concentration of partly destabilized and aggregation-prone states is decreased, thereby slowing misfolding. Additionally, the aggregation-blocking action of m^7^GDP may be also explained by a ligand-controlled equilibrium between the open and closed conformations of the protein^68^. Such stabilization could hinder the exposure of ‘amyloidogenic regions’ to the solution and the formation of new intramolecular interactions absent in the native structure.

Structural changes resulting from thermal denaturation were monitored using infrared (FT-IR) spectroscopy (**Figure 3B, E**). Prior to incubation at 37°C, the spectra indicated the dominance of α-helical structures, with the maximum of the amide I band at approximately 1647 cm^-1^ for hDcpS and 1651 cm^-1^ for CeDcpS. After 10 days of incubation at 37°C, the band shifted toward the lower wavenumber range (1630 cm^-1^ for hDcpS and 1631 cm^-1^ for CeDcpS), indicating the presence of parallel β-sheet structures. Of note, very similar IR signatures are observed for various other amyloid aggregates – e.g. of insulin-derived fibrils^69^.

The formation of the DcpS-derived amyloid-like fibrillar structures was visualized using transmission electron microscopy (TEM) (**Figure 3C, F**). Images were acquired after 10 days of incubation at 37 °C. Negative staining revealed that human DcpS protein can form fibrils which can be incorporated into thick bundles. These fibrils exhibited diameter of ∼8-9 nm. Similarly, *C.elegans* DcpS formed large bundles composed of fibrils with the diameter of ∼8-9 nm. The mature fibrils of DcpS appeared densely packed, hindering detailed morphological examination. To overcome this, the samples were sonicated before imaging. Notably, fibrils from both species were clearly visible and appeared predominantly linear, unbranched with noticeable bends. Additionally, oligomeric species were observed in the case of human DcpS samples.

### 4. DcpS^INS15^ mutation accelerates aggregation compared to the wild-type

In parallel to wild-type hDcpS, we also analyzed aggregation propensity of DcpS^INS15^ mutant^49^ (**Figure 4**). As it is shown in **Supplementary Figure 4A**, the far-UV CD spectra of DcpS^INS15^ were indistinguishable from those of WT DcpS, and the protein retained its ligand binding capacity and enzymatic activity (albeit at reduced level, ∼3x) (**Supplementary Figure 4B and 4F**). The T_agg_ (53°C) and apparent Tm (52°C) for DcpS^INS15^, based on the course of the nanoDSF thermograms, were significantly lower in comparison to the values obtained for WT hDcpS (around 4°C for T_agg_, and around 5°C for Tm) (**Supplementary Figure 4E**). This indicates that DcpS^INS15^ has much lower thermal protein stability and is more prone to aggregation at lower temperatures. Structural analysis based on far-UV CD spectroscopy further confirmed clear changes in the content of α-helices and β-sheets above 48°C, where the percentage of β-sheet began to increase (**Supplementary Figure 4D**). In an analogy to the unmutated form, the DcpS^INS15^ was not capable of refolding after cooling of the sample and subsequent incubation at 20°C (**Supplementary Figure 4C**). Analysis of the DcpS^INS15^ aggregation kinetics at 37°C using ThT assay showed a rapid increase in the fluorescence intensity upon incubation. The *T_50_* for DcpS^INS15^ was estimated to approximately 50 hours (at 0.5 and 0.25 mg/ml initial protein concentrations) and only to around 12 hours at 1mg/ml, what is comparable to *T_50_* value obtained for CeDcpS and four times faster than that of WT DcpS (*T_50_* ∼50 hours) (**Figure 4A**). ThT fluorescence intensity in in the presence of m^7^GDP was around 3 times weaker compared to the ligand-free sample, although around four times higher than that of wild type DcpS at the end-state. This clearly indicates that the mutation significantly accelerates aggregation of DcpS^INS15^. Visualization of DcpS^INS15^ amyloid-like fibrils by TEM revealed the presence of tightly associated fibril clusters and amorphous aggregates incorporating short fibrils (**Figure 4B**). These fibrils were ∼8-9 nm in diameter.

**Figure 4.**
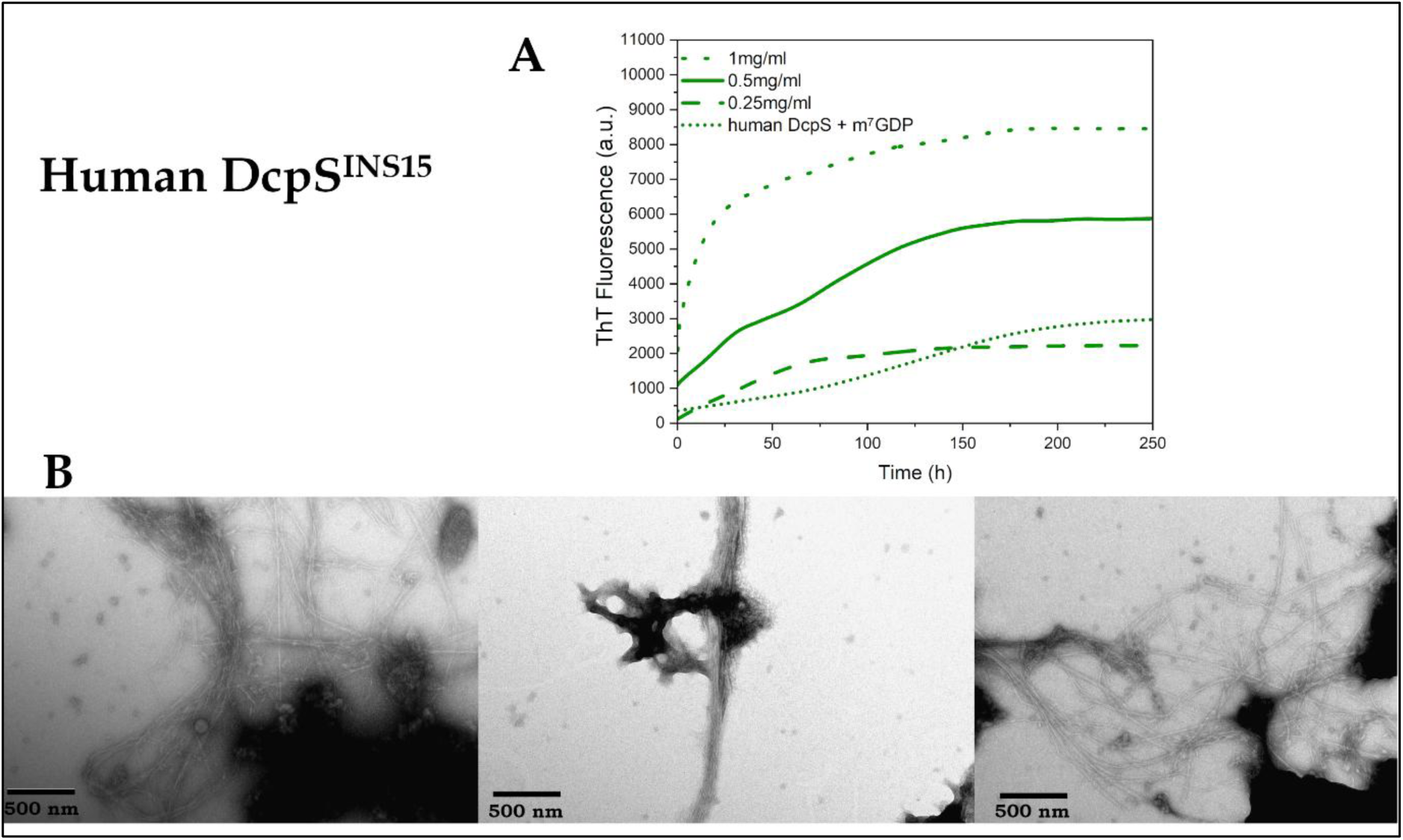
Amyloid-like fibril formation by human DcpS^INS15^ mutant protein. **A -** Kinetics of fibril formation by human DcpS^INS15^ analyzed by ThT assay. Experiments were performed at 37 °C for over 10 days of incubation for a range of DcpS concentrations: 0.25mg/ml (short dashed line), 0.5mg/ml (solid line) and 1mg/ml (dashed line). No detectable lag phase is seen regardless of protein concentration. Saturation of the ThT fluorescence signal and the end-state of fibril assembly is approached faster at higher protein concentrations used. Data of ThT assay in the presence of m^7^GDP ligand (at ligand:protein molar ratio 1:20), and at DcpS^INS15^ 0.5mg/ml concentration, is shown also (short dot line). Representative set of data of three independent experiments is shown. **B** – TEM images of DcpS^INS15^ fibrillar aggregates acquired after 10 days of incubation at 37°C. Samples were sonicated before uranyl acetate staining and TEM imaging. Three representative micrographs are shown. Scale bare represents 500 nm.

### 5. Droplet-like protein condensates of DcpS

The apparent lack (or shortening) of the lag (nucleation) phase in protein aggregation kinetics could be rationalized as symptoms of liquid-liquid phase separation and droplet formation preceding fibrillization^70,71^. Therefore, we chose to investigate the initial stage of the aggregation process of DcpS proteins and explore the possibility that they undergo aggregation/fibrillation via a liquid-liquid phase separation (LLPS) mechanism. Initial screening of databases, such as LLPSDB 2.0 and PhaSePro^72,73^, did not indicate DcpS as protein known to participate in LLPS formation *in vivo* or *in vitro*. To verify experimentally the hypothesis that DcpS undergoes LLPS, we analyzed microscopic images of DcpS in the presence of 5% and 10% concentrations of polyethylene glycol (PEG-4000) at a protein concentration of 1 mg/ml. PEG is used as the effective molecular crowding agent, mimicking the crowded intracellular environment, thereby facilitating the formation of protein-rich droplets^74^. As shown in the images obtained by confocal microscopy (**Figure 5**), no droplets were detected for any of the studied proteins at 0% PEG. However, at 5% PEG, distinct droplets were clearly seen, with diameters of approximately 1-3 μm for hDcpS and 2-5 μm for CeDcpS. In the case of CeDcpS, examples of two droplets in very close proximity can be found among the single droplets, indicating the potential fusion of condensates (marked by arrows, **Figure 5**). For the human DcpS mutant, only single spherical droplets with a diameter of 1-2 μm were detected and, in addition, particles of variable shape were also present, suggesting protein aggregation (**Figure 5**). At 10% PEG concentration, only irregular, probably aggregated structures were observed for each of the studied DcpS.

**Figure 5.**
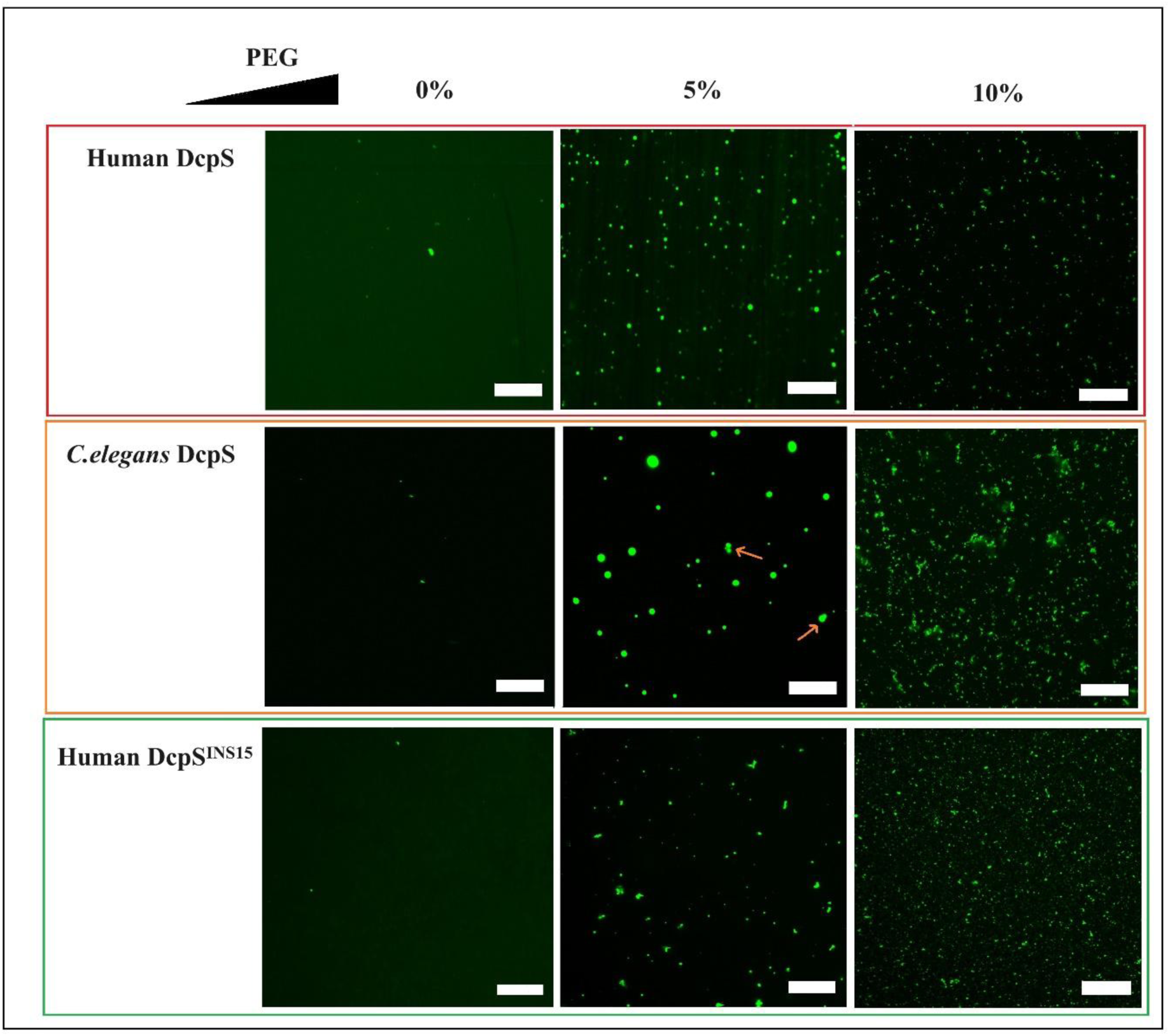
DcpS proteins undergo liquid-liquid phase separation in vitro. Confocal microscopic images of DcpS droplets visualized with Alexa Fluor 488 fluorescent dye, at protein: dye molar ratio 1:10 (incubation at room temperature for 1.5 hour). Images shows a distinct spherical droplets for *C.elegans* and human DcpS at 5% PEG-4000 crowding agent concentration. Potential fusion events between DcpS droplets are marked by red arrows. Data were obtained in a buffer containing 50mM HEPES, 150mM NaCl pH 7.5. Protein concentration 1mg/ml. Scale bars correspond to 20µm.

After 5 hours of incubation at 20°C, *C. elegans* and human DcpS droplets remained liquid-like and displayed greater dynamism, as reflected by the observation of a greater number of fusion events (**Supplementary Figure 5A**). However, after a shift to 37°C and 30 minutes of incubation, the CeDcpS droplets diverged into aggregate-like structures (**Supplementary Figure 5B**). In the case of human DcpS, droplets appeared smaller and not perfectly spherical, suggesting little fusion between condensates (**Supplementary Figure 5B**).

The presence of 5% PEG also had a positive effect on the aggregation kinetics of hDcpS, as the fluorescence signal of ThT started to increase earlier in comparison to the sample without PEG, with the *T_50_* estimated to ∼25 hours (compared to *T_50_* of PEG-free samples ∼80-100 hours) (**Supplementary Figure 6**). In the presence of 10% PEG, the ThT fluorescence remained “flat” over the incubation time, suggesting the presence of amorphic aggregates that are unable to form fibrillar structures. Taken together, these results demonstrate that DcpS proteins can undergo phase transition and form LLPS droplets *in vitro* in the artificially crowded environment, and that the formation of DcpS amyloid-like fibrils could occur via an LLPS-dependent pathway.

## Discussion

DcpS proteins have been extensively studied for their enzymatic activity in mRNA turnover, including their activity and specificity towards synthetic mRNA cap analogs - potent compounds used in designing stable therapeutic mRNA, such as mRNA vaccines, or as potential inhibitors of eIF4E—a protein overexpressed in various types of cancer^75–77^. Recently, the involvement of human DcpS and its mutants have been highlighted in the development of rare neurological disorders^49–51^ and in the regulation of neuronal development^53^, as well as in cancer diagnostics^46,47^.

The results of the present study demonstrate that DcpS can form protein amyloid-like fibrils. TEM images support such a statement, and visualized fibrillar aggregates possess morphology similar to well-known amyloid forming proteins^64,78,79^. Diameter of DcpS fibrils (∼8-9 nm) is similar to that of well-characterized amyloid fibrils of Aβ peptides (7-12 nm)^80,81^, observed in AD. Circular dichroism (CD) spectra indicated that thermally induced aggregation is accompanied by a transition of α-helical to β-sheet structures within the temperature range examined, with the resulting structural state capable of binding Thioflavin-T. Importantly, the increase in Thioflavin-T fluorescence was observed at the constant temperature (37°C), indicating the formation of amyloid-rich β-sheet aggregates by DcpS proteins from *C.elegans* and *H.sapiens* (physiological temperature for human). In the context of the results obtained, we included the human DpcS^INS15^ in the study - the first clinically significant DcpS mutation revealed. DpcS^INS15^ undergoes denaturation at lower temperatures and aggregates irreversibly into amyloid structures significantly faster than the wild-type DcpS, despite the far-UV CD spectra do not reveal any significant changes in the secondary structure. From the structural point, the DpcS^INS15^ is enriched in the additional aggregation-prone segment found in this additional 15-amino acid insertion, and molecular modeling revealed that this insertion is exposed to solvent^49^, what likely increases aggregate formation and decreases protein stability.

An intriguing aspect of the aggregation kinetics observed in all the DcpS species studied was the shape of the aggregation curve, which did not correspond to the classical sigmoidal pattern associated with the nucleation phase of fibril formation^21^. While factors such as temperature or salt concentration may contribute to the apparent absence of a lag phase^82^, it is also possible that rapid formation of new aggregates occurs due to pre-existing amyloid fibrils or other seeding agents^19^. They accelerate fibril growth by providing a catalytic platform for further elongation (secondary seeding), as observed for lysozyme fibrils^78^, Aβ fibrils of Alzheimer’s disease^83^, or α-Synuclein amyloid aggregates associated with Parkinson’s disease^84^. Cross-seeding - where amyloid structures of one protein serve as templates for the aggregation of another—is also a well-documented process that promotes fibril formation (e.g. in studies of the interaction between Tau and Aβ42 in Alzheimer’s disease or α-synuclein and Aβ in Parkinson’s disease^85^, or in recent reports suggesting that SARS-CoV-2 N-protein and S1 RBD protein may cross-seed α-synuclein aggregation, potentially increasing the risk of Parkinson’s disease post-transmission^86–88^. However, our results unequivocally demonstrate that fragments of sonicated mature fibrils of the wild-type hDcpS, or the DcpS^INS15^ mutant used for cross-seeding experiments, do not promote secondary nucleation or shorten the fibril growth phase of human DcpS (**Supplementary Figure 7**).

Numerous studies showed that protein liquid-liquid phase separation could trigger the formation of protein amyloid aggregates^70,71^, and it was also proposed that LLPS could act as an alternative nucleation pathway that accelerates the aggregation^89^. In accordance with observations for other proteins that form droplets^70,90^, it is possible that the absence of a discernible lag phase for *C. elegans*, human DcpS, and its mutant is due to the existence of a liquid-liquid phase separation (LLPS) mechanism. Indeed, the microscopy imaging results showed that both human and *C. elegans* DcpS proteins can form droplets under molecularly crowded conditions at physiological pH and salt concentration. Over time, two or more droplets can merge to form larger droplets^91^, and the frequency and extent of fusion events may increase during incubation time, leading to the formation of larger droplets - a phenomenon referred to as maturation^92^. Similar to the reported examples^91,92^, we observed such fusion events of LLPS droplets also for human and *C. elegans* DcpS, further supporting the notion that DcpS proteins undergo a liquid-liquid phase separation process. Furthermore, the lack of changes in aggregation rate for DcpS proteins depending on concentration can also be attributed to LLPS^90^. During the early stages of aggregation kinetics measurements, the detection of Thioflavin T fluorescence signal above zero suggests conformational changes involving β-sheet transitions in condensates^70^.

Available studies suggest that the proteins with the intrinsically disordered regions or low complexity domains tend to have a higher propensity to form droplets^93,94^. However there are also examples of a few globular proteins undergoing liquid-liquid separation^95–97^, and now DcpS can be added to this list. The intriguing question is why DcpS proteins are not abundant in membraneless organelles like P-bodies or stress granules involved in RNA processing or mRNA degradation, similarly to other RNA-interacting proteins^27,98^. One assumption is that LLPS is known to be facilitated by long RNA molecules^99,100^, whereas DcpS processes short mRNA transcripts up to 10 nucleotides, but with the highest effectivity towards dinucleotides^42^. However, there are known droplet-forming proteins that do not interact with RNA^101,102^, therefore DcpS proteins may also have this property, which in turn could be connected to their biological functions beyond mRNA degradation (although these functions are currently unknown).

Our study has also shown that the binding of the m^7^GDP cap analog by the DcpS protein significantly limits its aggregation process. As such, m^7^GDP and other specific ligands of DcpS could be tested as potential inhibitors of DcpS-derived fibril formation or used as a starting point in designing of novel anti-aggregation compounds, similarly to the creation of compound libraries with therapeutic significance that block protein aggregation^103,104^.

Given that proteins forming amyloids *in vitro* do not always do so *in vivo*^105^, and that only about 33% of proteins that could form LLPS droplets *in vitro* undergo this process within the range of their cellular concentrations (with regard to the PaxDb database)^106^, a relevant question is whether DcpS forms functional (or pathological) fibrils or biomolecular condensates *in vivo*. And whether DcpS droplets could then serve as, for example, an intermediate phase on the pathway to the formation of amyloid-like fibrils, or potential physiological mechanism connected to regulation of its enzymatic activity^107^. This requires further investigation, in particular the visualization of this process in the cellular environment under different conditions. Especially, such an analysis should focus on the human mutant protein with a 15 amino acid insertion, as well as other known mutants associated with Al-Rhad syndrome^49–52^.

As DcpS proteins are members of the histidine triad (HIT) superfamily of enzymes, it raises a question whether any other HIT protein has the ability to form fibrillar structures in vitro (and in vivo). Although not amyloid-like fibrils, the recent report showed that HINT1 protein from this superfamily is turned out into long polymers (from 2 to ∼10 000 subunits) and that this is promoted by the presence of diadenosine tetraphosphate Ap4A^108^. This HINT1 polymerization is enhanced by higher concentrations of this dinucleotide, leading to filaments of ∼10μm in length (and 10-250 nm in width).

In conclusion, our results clearly showed that DcpS scavenger proteins, of human and *C.elegans* nematode origin, form amyloid fibrils through the LLPS-assisted pathway. The clinically significant DpcS^INS15^ mutant is significantly less thermally stable and tends to aggregate into amyloid structures significantly faster than the wild-type DcpS.

## Methods

### Recombinant Protein Production and Purification

Human and *C.elegans* DcpS were expressed in the *Eschericha coli* Rosetta2 (DE3) from the pET30a(+)hDcpS or pET30a(+)CeDcpS vectors, according to the procedure described previously^54^. Briefly, protein expression was induced with 0.4 mM IPTG at OD600 around 0.5-0.8, and the bacterial culture was incubated in LB medium at 18°C overnight. The cell pellet collected by centrifugation (7000g, 10 min) was resuspended in the ice-cold lysis buffer (50 mM TRIS pH 7.5, 300 mM NaCl, 1% Triton X-100 and 20 mM imidazole), sonicated and centrifuged at 4°C, 30000g, for 3 hours. The supernatant was applied to a HisTrapHP column (Cytiva) equilibrated with 50 mM TRIS pH 7.5, 300mM NaCl and the DcpS protein was eluted with an imidazole gradient (20-600 mM) with flow rate 0.5ml/min. The final purification step was gel filtration on a Superdex 200 10/300GL column (GE Healthcare Life Sciences) using ÄKTA FPLC protein purification system (GE Healthcare Life Sciences) and appropriate (for performed later experiments) elution buffer: 50mM TRIS pH 7.5, 150mM NaCl, 0.2mM TCEP, or 15 mM TRIS pH 7.5, 80 mM NaClO_4_, and 0.2 mM TCEP. The purified protein was stored at 4°C.

Human DcpS^INS15^ was purified according to the above procedure using the pET30a+INS15hDcpS plasmid, which was generated by replacing the wild-type DcpS open reading frame with the cDNA encoding a 15-amino acid insertion^49^ at the NdeI/BamHI restriction sites of the pET30a+ vector (BioCat GmbH, Germany).

### Nano-differential scanning fluorimetry (nanoDSF)

NanoDSF experiments for DcpS were performed using a NanoDSF Prometheus NT.48 device equipped with backreflection mode (NanoTemper Technologies, München, Germany). Upon measurement, capillaries were filled with a DcpS protein suspension at a final concentration of 0.5 mg/mL in 50 mM TRIS, 150mM NaCl, 0.2mM TCEP, pH 7.5 buffer and then subjected to thermal stress from 20°C to 80 °C, with a ramp rate of 0.5°C/min. Fluorescence emission was recorded at 330 nm and 350 nm following UV excitation at 280 nm. Protein aggregation was monitored simultaneously using back-reflection optics. Thermal stability parameters, including Tm and the onset temperature of aggregation (Tagg), were determined by implemented PR.ThermControl v2.1.2 software (NanoTemper Technologies).

### Circular dichroism (CD) spectroscopy

Circular dichroism (CD) spectra of DcpS were recorded in 15 mM TRIS, 80 mM NaClO₄, and 0.2 mM TCEP (pH 7.5) buffer with a final enzyme concentration of 0.2 mg/mL using a CD Chirascan Plus spectrometer (Applied Photophysics Ltd., Leatherhead, UK). Conformational changes in the protein secondary structure were monitored in the far-UV range between 190 nm and 260 nm, at 0.5 nm intervals, in a quartz cuvette with a path length of 0.5 mm. The far-UV CD spectra were recorded at temperatures range from 20°C up to 64°C (for human DcpS) or up to 52°C (for *C.elegans* DcpS). The temperature was increased stepwise from 20°C at a heating rate of 1°C/0.5min. The percentage of protein secondary structures was determined from the recorded spectra using BeStSel software^60^.

In renaturation experiments, far-UV CD spectra were recorded at 20°C after human DcpS or *C. elegans* DcpS being heated to 56 °C or 44 °C, respectively (at a heating rate of 1°C/0.5min), followed by gradual cooling to 20°C and incubation for 12 hours (before measurement).

### Differential scanning calorimetry (DSC)

Differential scanning calorimetry (DSC) measurements were carried out using a MicroCal VP-DSC calorimeter (Malvern Instruments, UK) in the temperature range of 20–100 °C, with a scanning rate of 0.5 °C per minute. The protein sample was prepared at a concentration of 0.7 mg/mL in a buffer containing 50 mM TRIS, 150 mM NaCl, and 0.2 mM TCEP at pH 7.5. Prior to DSC measurements, both the protein solution and the buffer used as a reference were degassed under vacuum. For each experiment, scans buffer–buffer and sample–buffer were performed under identical conditions. Final thermograms were obtained by subtracting the appropriate reference baseline (buffer–buffer) from the protein thermogram. The resulting excess heat capacity curves were analyzed using the MicroCal LLC DSC plug-in for Origin 7.0 software, provided with the instrument.

### Thioflavin T (ThT) fluorescence assay

DcpS protein samples at concentrations of 0.25 mg/ml, 0.5 mg/ml, and 1 mg/ml (in a 50 mM TRIS, 150 mM NaCl, and 0.2 mM TCEP pH 7.5 buffer) were incubated in the presence of thioflavin T (Sigma-Aldrich) at a final concentration of 10 μM (0.0031886 mg/ml) in a total volume of 200 μl.

Each sample was pipetted into a 96-well transparent plate (Greiner) and sealed with adhesive seals (Biorad, Microseal ‘B’Adhesive Seals). ThT fluorescence was monitored using Synergy H1 (BioTek) microplate reader, at 487 nm emission and 440 nm excitation wavelengths. Measurements were performed at 37°C with orbital agitation at 200 rpm over 10 days.

### Formation of DcpS-derived Amyloid Fibrils

To form amyloid fibrils, DcpS at 0.5 mg/mL concentration (in a 50 mM TRIS, 150 mM NaCl, and 0.2 mM TCEP pH 7.5 buffer) in 200 μL volume were incubated at 37°C over 10 days with continuous shaking in a microplate reader (Synergy H1, BioTek). Fibril formation was monitored in a parallel sample well where thioflavin T was present (ThT fluorescence assay). After 10 days samples from wells without ThT were collected, and the morphology of the resulting DcpS-derived fibrils was examined by transmission electron microscopy (TEM). Samples of CeDcpS- and DcpS^INS1^-derived fibrils were sonicated (for 3 min in Elmasonic S30H ultrasound sonicator) prior to TEM imaging.

### Transmission Electron Microscopy

DcpS-derived fibrils, obtained as described above, were applied to a 400-mesh copper grid, coated with Formvar and carbon (Ted Pella), for 40 seconds and subsequently negatively stained with a 2% (w/v) water solution of uranyl acetate (Serva Electrophoresis GmbH) for 25 seconds. The grids were dried at room temperature and the micrographs were collected using JEM 1400 transmission electron microscope (JEOL Co., Japan) equipped with 11-megapixel MORADA G2 TEM camera (EMSIS GmbH, Germany), operated at 80 kV.

### FT-IR Measurements

Infrared spectra were collected using a Nicolet iS50 FTIR spectrometer (Thermo, USA) equipped with a single-reflection diamond ATR (Attenuated Total Reflectance) accessory and a DTGS detector. Centrifuged samples of DcpS aggregates collected from the plate reader (after 10 days incubation at 37°C) were washed several times with H_2_O to remove buffer components. Liquid samples approximately 10 µL in volume were transferred onto the diamond surface of the ATR accessory and gently dried in situ. Subsequently, infrared spectra of the resulting films were recorded. Typically, 32 interferograms were collected at a resolution of 2 cm^-^¹ to produce a single spectrum. The spectral data were processed with GRAMS software (Thermo).

### Fluorescence Microscopy Imaging

Liquid–liquid phase separation (LLPS) of protein samples was monitored using confocal fluorescence microscopy. Proteins were fluorescently labeled with ATTO 488 dye (Sigma-Aldrich) according to the manufacturer’s protocol. The DcpS protein was prepared at a final concentration of 1 mg/mL in 50 mM HEPES, 150 mM NaCl, pH 7.5, and incubated with ATTO 488 at room temperature for 1.5 hours in a final volume of 50 μl. The molar ratio of protein to dye during labeling was 10:1. Unbound dye was removed using Amicon Ultra-15 centrifugal filters (10 kDa MWCO, Merck).

To induce droplet formation, labeled protein samples were incubated under the same buffer conditions in the presence of 5% or 10% PEG-4000 (as a macromolecular crowding agent). For imaging, 5–10 μl of each sample was applied to a glass slide and imaged either immediately or after the indicated incubation time. Fluorescence images were acquired using a Zeiss LSM 700 confocal microscope equipped with a 40× oil immersion objective and processed using ZEISS ZEN 3.3 (Blue Edition) software.

## Acknowledgements

We would like to thank Professor Krzysztof Nieznański for the insightful discussion and valuable feedback, which significantly improved this manuscript. This work was supported by grant from the Polish National Science Centre, UMO/2017/27/B/NZ1/01859. TEM observation was performed in the Laboratory of Electron Microscopy, Nencki Institute of Experimental Biology, PAS, Warsaw, Poland, using infrastructure supported by EuBI Polish Node “Advanced Light Microscopy Node Poland”. This work is dedicated to Professor Edward Darzynkiewicz, who passed away during the writing of this manuscript.

