## Supplementary Material for "Liquid-Liquid Phase Separation-mediated formation of amyloid fibrils from DcpS scavenger enzymes"

**Supplementary Table 1.** Aggregation prone regions (APR) predicted by WALTZ algorithm for human DcpS, mutant human DcpS with 15aa insertion and *C.elegans* DcpS. APR segments are numbered according to appearance in the amino acid sequence of DcpS.

| APR | **Human DcpS** | **Human DcpS $^{INS15}$** | ***C.elegans* DcpS** |
|---|---|---|---|
| segment no.1 | $^{173}$IQWVYNI$^{179}$ | $^{173}$IQWVYNI$^{179}$ | $^{147}$LNWVYN$^{152}$ |
| segment no. 2 | $^{199}$GFVLIP$^{204}$ | $^{199}$GFVLIP$^{204}$ | $^{187}$LENLYVLAI$^{195}$ |
| | | $^{218}$WNVLIS$^{223}$ | |

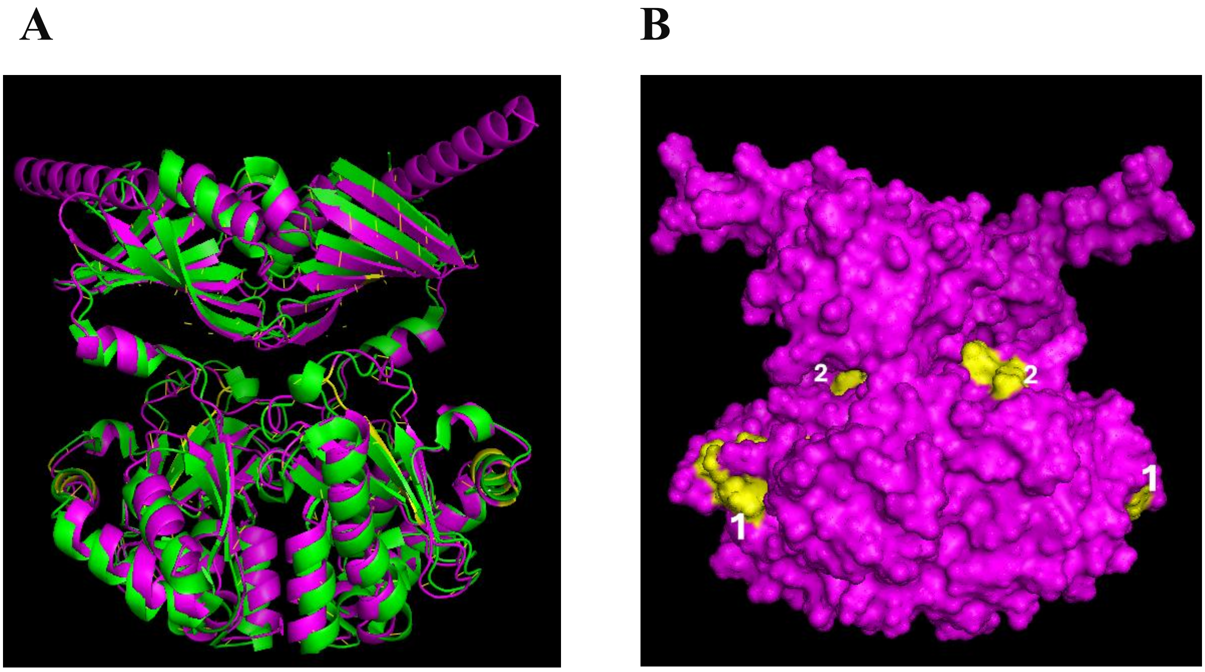


**Supplementary Figure 1. (A)** Comparison of AlphaFold prediction model of *C.elegans* DcpS structure (**magenta**) with human DcpS structure (PDB:1XML) (**green**). *C.elegans* DcpS structure was predicted using AlphaFold 2 (Jumper *et.al.,* 2021). **(B)** Localization of the predicted APR segments (no.1 and no.2) (**yellow**) within the predicted dimeric structure of *C.elegans* DcpS.

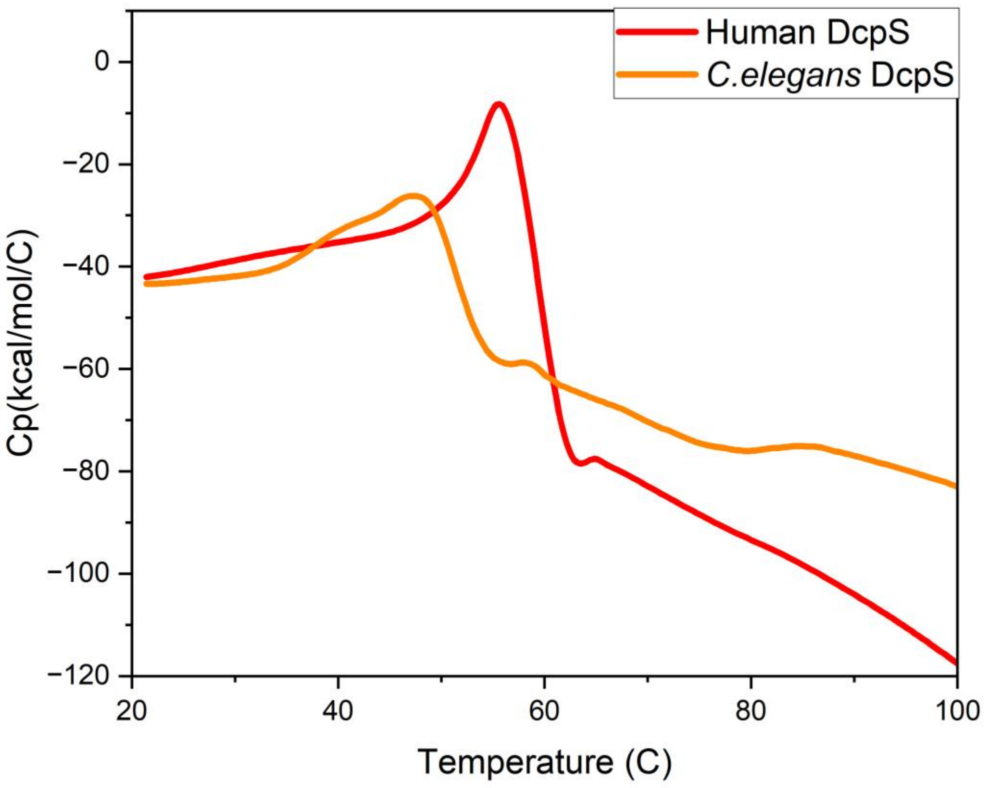


**Supplementary Figure 2.** DSC heat capacity profiles demonstrating changes in the apparent molar heat capacity of aggregating DcpS proteins from humans and *C. elegans*. A positive peak is observed, indicating an endothermic transition at 57°C for human DcpS and 48°C for *C. elegans* DcpS, preceding another endothermic transition. Experiments were performed at 0.7 mg/ml protein concentration in 50 mM TRIS, 150 mM NaCl, 0.2 mM TCEP, pH 7.5, at a scanning rate 0.5°C/min.

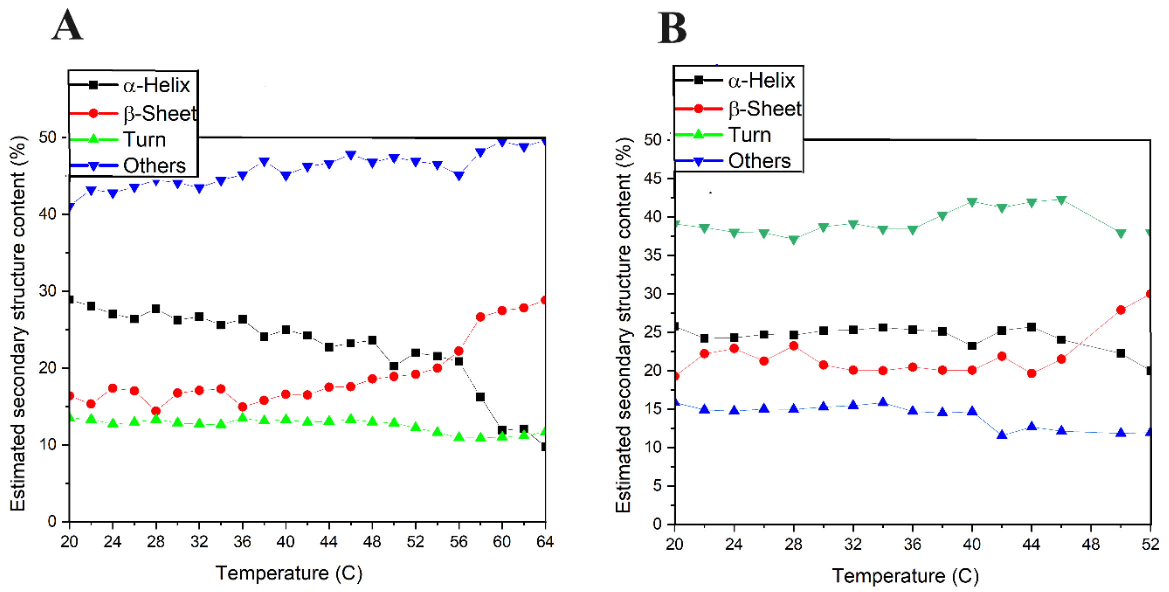


**Supplementary Figure 3.** Estimated secondary structure content of human **(A)** and *C. elegans* **(B)** DcpS by BeStSel server according to CD measurements at temperatures ranging from 20°C to 64°C for human DcpS and form 20°C to 52°C for *C. elegans* DcpS. The percentages of α-helix and β-sheet are represented by filled black squares and red circles, respectively.

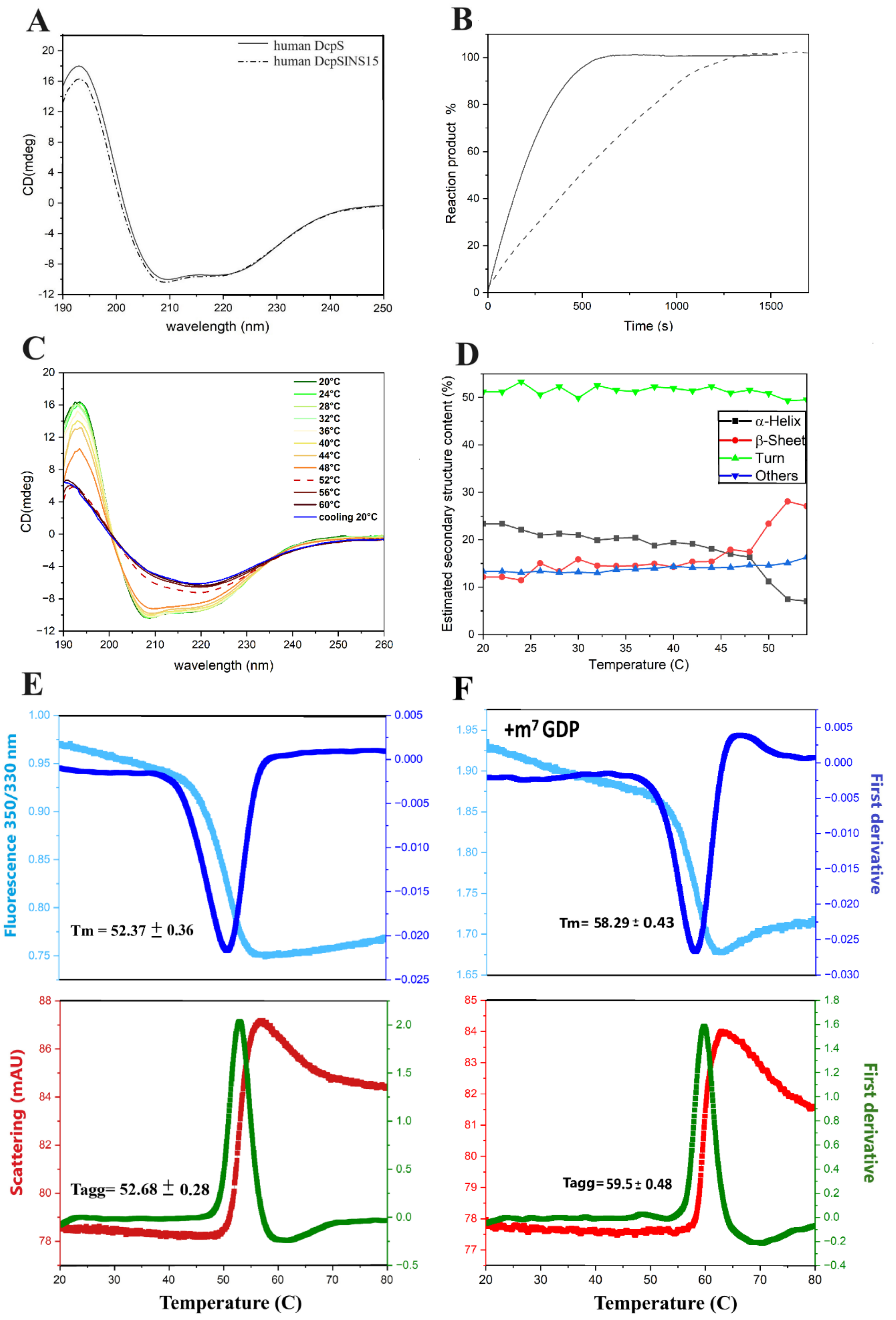


**Supplementary Figure 4. Biophysical properties of human mutant DcpS$^{INS15}$ protein.**

**Supplementary Figure 4. Biophysical properties of human mutant DcpS$^{INS15}$ protein**. (continued)

**A**. Comparative far-UV CD spectra of human DcpS and its insertional mutant DcpS$^{INS15}$

**B.** Kinetics of cap hydrolysis catalyzed by human DcpS (smooth line for WT DcpS and dashed line for DcpS$^{INS15}$). Reaction progress curves were obtained for the 10 μM m$^7$GpppG dinucleotide substrate in the presence of 30 nM of the indicated enzyme.

**C**. Overlaid smoothed far-UV CD spectra of DcpS$^{INS15}$ obtained at a temperatures ranging from 20°C to 60°C (what is 8°C above the estimated with nanoDSF melting temperature Tm, dashed line). Green line is for the spectrum obtained at 20°C, followed by the series of rainbow-colored lines corresponding to plots obtained at 2°C intervals. In dark blue is shown spectrum obtained after cooling at 20°C for 12 hours the DcpS$^{INS15}$ protein sample heated to 52°C. Spectra were acquired at the heating rate 0.5°C/min.

**D**. Estimated secondary structure content by BeStSel server according to CD measurements at temperatures ranging from 20°C to 60°C. The percentages of α-helix and β-sheet are represented by filled black squares and red circles, respectively.

**E** and **F.** Thermal stability of DcpS$^{INS15}$ in the absence (**E**) or presence of m$^7$GDP nucleotide ligand (**F**), measured by nanoDSF (upper panels). Presented thermograms display the first derivative (dark blue) of fluorescence signal intensity ratio at 350nm and 330nm emission wavelength (light blue) under thermal unfolding in the temperature range 20°C – 80°C. The temperature at which melting transition (Tm) occur is shown.

Bottom panels in **E** and **F** shows measured scattering profiles for DcpS$^{INS15}$ (red line) in temperature range 20°C – 80°C (using NanoDSF Prometheus NT.48 with back-reflection optics). Green line corresponds to the first derivative of a scattering profile, and calculated aggregation temperature is shown. For nanoDSF thermal stability and scattering measurments the heating rate was 0.5°C/min.

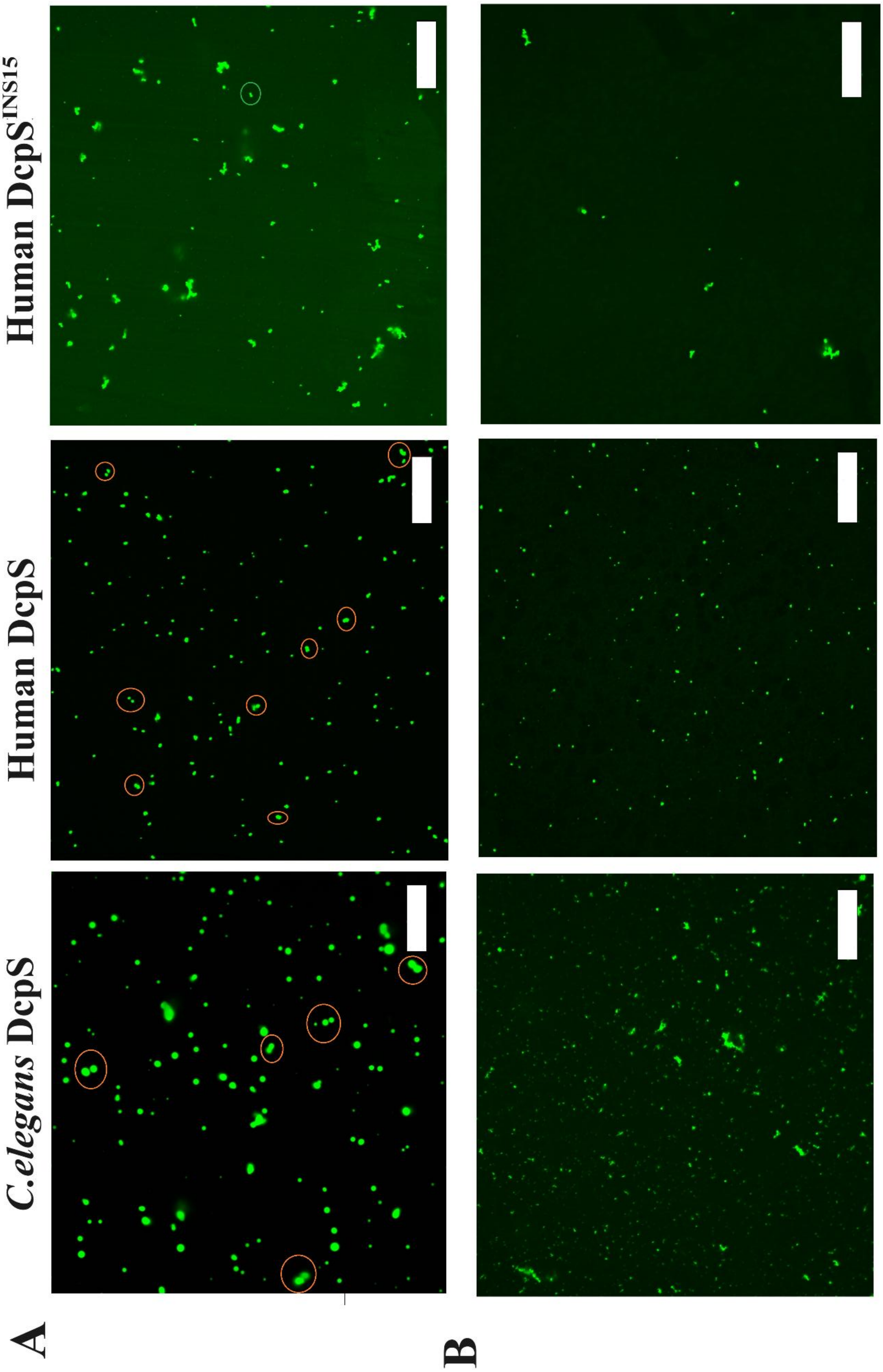


**Supplementary Figure 5.**

**Supplementary Figure 5. Confocal microscopic images of DcpS droplets.** (continued)

(**A)** 5 hours incubation in 20°C in presence of 5% PEG 4000 result in fusion event for *C.elegans* DcpS and human DcpS, while mutant DcpS$^{INS15}$ presents irregular probably aggregated structures. Representation of potential fusion events between DcpS droplets are marked by circles.

(**B)** After 30 minutes incubation in 37°C, small potential aggregates appeared for *C.elegans* DcpS and human mutant DcpS$^{INS15}$, and human WT DcpS droplets were smaller with weaker tendency to fusion than in 20°C. Data obtained in a buffer containing 50mM HEPES, 150mM NaCl pH 7.5. Scale bars 20µm.

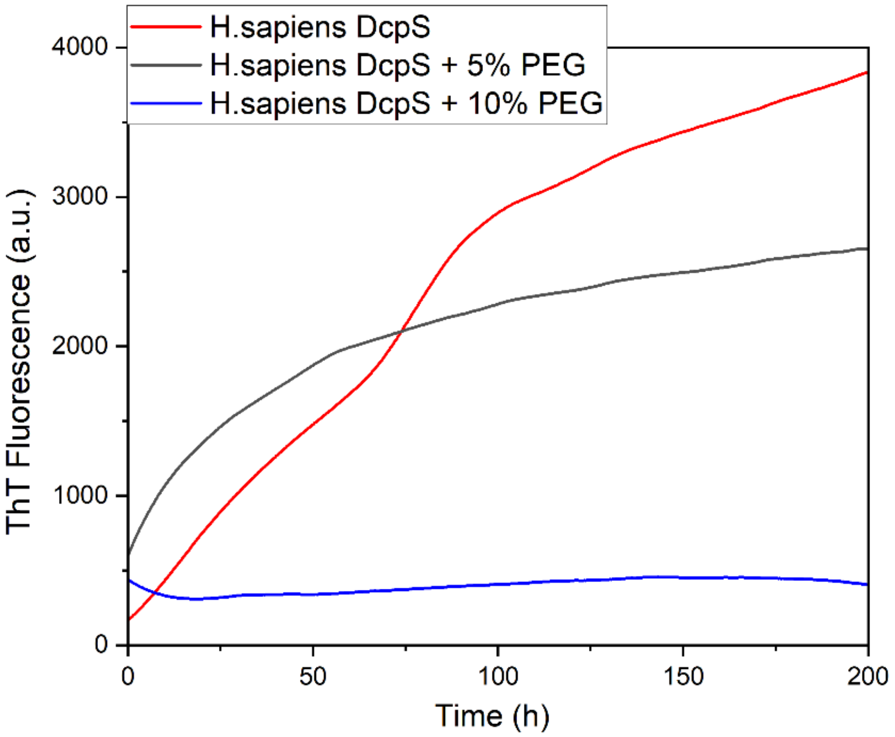


**Supplementary Figure 6.** Thioflavin T fluorescence intensity of human DcpS under molecular crowding conditions: without PEG 4000 (red line - control), with 5% PEG 4000 (grey line) and 10% PEG 4000 (blue line). Protein concentration – 0.5mg/ml.

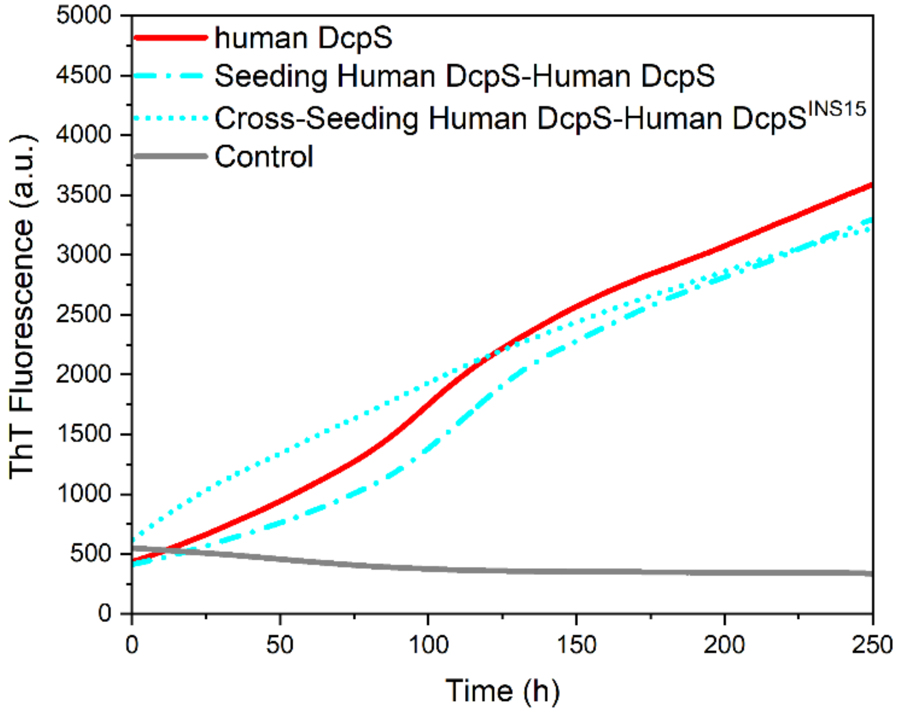


**Supplementary Figure 7.** Aggregation kinetics by ThT assay of human DcpS in the presence or absence of homogenic or heterogenic preformed DcpS seeds. Samples of purified DcpS (0.5mg/ml) were incubated alone (solid red line) or with sonicated fibrillar aggregates of wild-type human DcpS (dash-dot cyan line) or its mutant - DcpS$^{INS15}$ (dotted cyan line), at seeds to DcpS ratio 20:1. Grey solid line correspond to control (in 50mM TRIS, 150mM NaCl, 0.2mM TCEP, pH 7.5 buffer) with sonicated human DcpS fibrillar aggregates to make a sure that seeds do not generate false positive signal at concentrations used in seeding experiments.
